# ER-positive breast cancer cells are poised for RET-mediated endocrine resistance

**DOI:** 10.1101/098848

**Authors:** Sachi Horibata, Edward J. Rice, Hui Zheng, Lynne J. Anguish, Scott A. Coonrod, Charles G. Danko

## Abstract

The RET tyrosine kinase signaling pathway is involved in the development of endocrine resistant ER+ breast cancer. However, the expression of the RET receptor itself has not been directly linked to clinical cases of resistance, suggesting that additional factors are involved. We show that both ER+ endocrine resistant and sensitive breast cancers have functional RET tyrosine kinase signaling pathway, but that endocrine sensitive breast cancer cells lack RET ligands that are necessary to drive endocrine resistance. Transcription of one RET ligand, GDNF, is necessary and sufficient to confer resistance in the ER+ MCF-7 cell line. In patients, RET ligand expression predicts responsiveness to endocrine therapies and correlates with survival. Collectively, our findings show that ER+ tumor cells are “poised” for RET mediated endocrine resistance, expressing all components of the RET signaling pathway, but endocrine sensitive cells lack high expression of RET ligands that are necessary to initiate the resistance phenotype.

## Introduction

Estrogen receptor alpha (ERα) is the major driver of ∼75% of all breast cancers. Current therapies for patients with ER+ breast cancer are largely aimed at blocking the ERα signaling pathway. For example, tamoxifen blocks ERα function by competitively inhibiting E2/ERα interactions^1^ and fulvestrant promotes ubiquitin-mediated degradation of ERα^2^. Endocrine therapies are estimated to have reduced breast cancer mortality by 25-30%^3^. However, despite the widespread success of endocrine therapies, approximately 40-50% of breast cancer patients will either present with endocrine-resistant breast cancer at the time of diagnosis or progress into endocrine-resistant disease during the course of treatment^4^. Thus, there remains an urgent need to further elucidate the mechanism of endocrine resistance.

Numerous studies have now identified growth factor-stimulated signaling “escape” pathways that may provide mechanisms for cell growth and survival that are independent of E2. Foremost among these, the RET tyrosine kinase signaling pathway has been associated with endocrine resistance in both cell culture models as well as in primary tissues^5–8^. These studies have led to effective new biomarkers based on the downstream targets of RET signaling^6^. However, resistance by the RET signaling pathway has proven complex, relying in some cases of a functional ERα to drive resistance in aromatase inhibitor models^6^. Furthermore, genetic alterations in RET or its co-receptor, GFRA1, do not appear to be common in clinical cases, suggesting that additional factors are involved. A better understanding of the transcriptional targets of RET-mediated signaling pathways as well as understanding how these pathways crosstalk with ERα signaling will likely aid in the development of new predictive biomarkers and new targets for therapeutic intervention.

Here, we used Precision Run-On and Sequencing (PRO-seq) to comprehensively map RNA polymerase in tamoxifen-sensitive (TamS) and resistant (TamR) MCF-7 cells^9^. This approach is highly sensitive to immediate and transient transcriptional responses to stimuli, allowing the discovery of target genes within minutes of activation [ref 12-16]. Moreover, active transcriptional regulatory elements (TREs) can be detected by this method, including both promoters and distal enhancers, as these elements display distinctive patterns of transcription that can aid in their identification^10–15^. Among the 527 genes and 1,452 TREs that differ in TamS and TamR MCF-7 cells, we identified glial cell line-derived neurotrophic factor (GDNF), a ligand of RET tyrosine kinase receptor, to be upregulated in TamR MCF-7 cells. Remarkably, we found that all of the proteins necessary to drive endocrine resistance through RET receptor signaling are expressed in TamS MCF-7 cells, with the exception of a single limiting protein, GDNF or any of the other RET ligands (GDNF, NRTN, ARTN, or PSPN). To test this model, we manipulated GDNF expression in MCF-7 cells and found that high GDNF expression is both necessary and sufficient for tamoxifen resistance in our MCF-7 cell model. Several lines of evidence suggest that RET ligands are the limiting reagent in clinical samples as well, including ample expression of RET and its co-receptors, but limiting expression of GDNF and the other RET ligands in primary tumors. Additionally, RET ligand expression is predictive of responsiveness to endocrine therapies in breast cancer patients. Taken together, our studies support a model in which tamoxifen sensitive and resistant cells are ‘poised’ for RET-mediated endocrine resistance by expressing RET and its co-receptor, but are limited by the abundance of RET ligands to drive a resistant phenotype.

## Results

### Transcriptional differences between endocrine sensitive and resistant MCF-7 cells

Although MCF-7 cells are ER+ and usually require E2 for growth and proliferation, a subset of the heterogeneous MCF-7 cell population continues to grow in the presence of anti-estrogens such as tamoxifen^9,16^. We hypothesized that the resistant cells display a unique transcriptional program which can be used to identify factors that play a causative role in tamoxifen resistance. We used PRO-seq to map the location and orientation of RNA polymerase in two tamoxifen sensitive (TamS) and two *de novo* resistant (TamR) MCF-7 cell lines that were clonally derived from parental MCF-7 cells^9^. Consistent with the Gonzalez-Malerva study, we found that the TamS lines (TamS; B7^TamS^ and C11^TamS^) were sensitive to as little as 1 nM of tamoxifen, while the TamR lines (TamR; G11^TamR^ and H9^TamR^) were not affected at concentrations as high as 100 nM (Fig. 1a). PRO-seq libraries were prepared from all four cell lines (Fig. 1b), as previously described^17,18^, and sequenced to a combined depth of 87 million uniquely mapped reads (Supplementary Table 1). We quantified the similarity of transcription in the MCF-7 cell subclones by comparing the Pol II abundance in annotated gene bodies. Unbiased hierarchical clustering grouped B7^TamS^ and C11^TamS^ TamS lines into a cluster and left G11^TamR^ and H9^TamR^ TamR lines as more distantly related outgroups (Fig. 1c). Although TamR cells clustered independently, all four MCF-7 clones are nevertheless remarkably highly correlated (Spearman’s Rho > 0.95), suggesting that relatively few transcriptional changes are necessary to produce the tamoxifen resistance phenotype.

**Figure 1.**
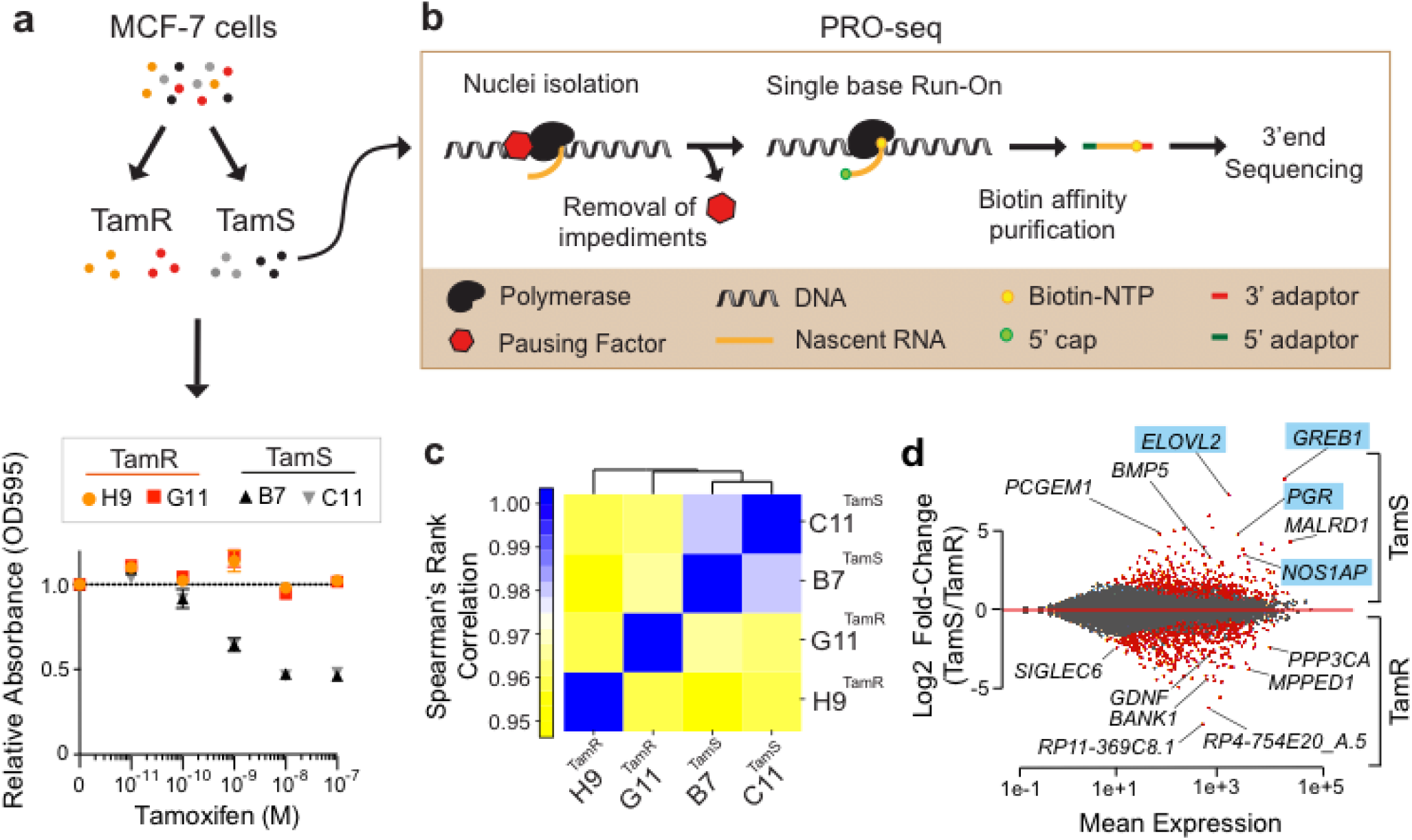
PRO-seq provides a genome-wide location of active RNA polymerase. (**a**) Cell viability of tamoxifen sensitive (TamS; B7^TamS^ and C11^TamS^) and resistant (TamR; G11^TamR^ and H9^TamR^) MCF-7 cells upon treatment with 0 (vehicle; EtOH), 10^-11^, 10^-10^, 10^-9^, 10^-8^, or 10^-7^ M of tamoxifen for 4 days. Data are represented as mean ± SEM (n=3). (**b**) Experimental setup for PRO-seq. PRO-seq libraries were prepared from all four cell lines grown in the absence of tamoxifen for 3 days. (**c**) Spearman’s rank correlation of RNA polymerase abundance in the gene bodies (+1000 bp to the annotation end) of TamS and TamR cell lines. (**d**) MA plot showing significantly changed genes (red) that are higher in TamS (top) or TamR (bottom) MCF-7 lines. Genes highlighted in the plots which are ERα targets are highlighted in blue.

We identified 527 genes that are differentially transcribed in TamS and TamR MCF-7 cells (1% FDR, DESeq2^19^), 341 of which were transcribed more highly in TamS and 186 more highly in TamR cell lines (Fig. 1d). Several of the differentially transcribed genes, including, for example, *PGR*, *GREB1*, *IGFBP5*, *HOXD13*, and *GDNF,* were identified in other models of endocrine resistance^6,7,20–23^, supporting our hypothesis that transcriptional changes in the MCF-7 model are informative about endocrine resistance.

### ER target genes are uniquely expressed in tamoxifen-sensitive MCF-7 cells

To confirm that transcriptional changes detected using PRO-seq lead to differences in mRNA abundance, we validated transcriptional changes in *PGR* (Fig. 2a) and *GREB1* (Fig. 2b) between the B7^TamS^ and G11^TamR^ MCF-7 cells using qPCR (Fig. 2c and 2d). Many of the differentially transcribed genes are targets of ERα signaling, including *PGR, GREB1*, *NOS1AP*, and *ELOVL2*, (Fig. 1d) suggesting that changes between TamR and TamS MCF-7 cells can be explained in part by differences in the genomic actions of ERα. To test for an enrichment of ERα target genes, we used an independent GRO-seq dataset^24^ to investigate whether immediate transcriptional changes following E2 treatment are correlated with genome-wide changes in TamS and TamR MCF-7 cells. We found that genes up-regulated by 40 minutes of E2 treatment tend to be transcribed more highly in TamS MCF-7 cells, and genes down-regulated by E2 are more highly transcribed in TamR cell lines (Fig. 2e). Thus, our data demonstrate global changes in the genomic actions of ERα in tamoxifen resistance in this MCF-7 model system.

**Figure 2.**
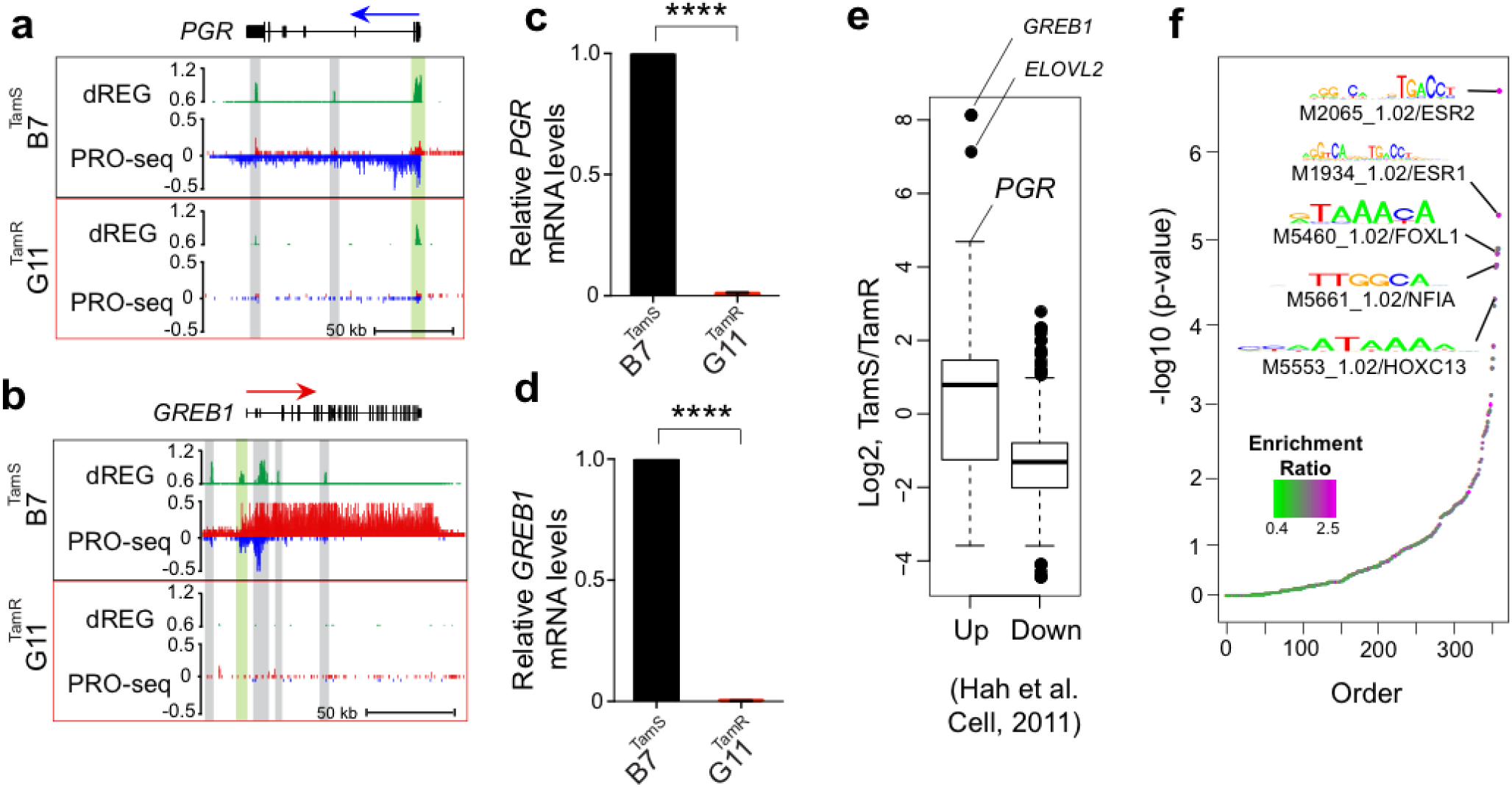
ER target genes are uniquely expressed in TamS cells. (**a-b**) Transcription near the *PGR* (**a**) and *GREB1* (**b**) loci in B7^TamS^ and G11^TamR^ cells. PRO-seq densities on the sense and anti-sense strand are shown in red and blue, respectively. dREG scores are shown in green. Enhancers and promoters are shown in grey and light green shading, respectively. Arrows indicate the direction of gene annotations. (**c-d**) *PGR* (**c**) and *GREB1* (**d**) mRNA expression levels in B7^TamS^ and G11^TamR^ cells. Data are represented as mean ± SEM (n=3 for *PGR*; n = 4 for *GREB1*). **** p < 0.0001. G11^TamR^ is normalized to B7^TamS^. (**e**) Boxplots represent fold-change between TamS and TamR of genes that are either up- or down-regulated following 40 minutes of estrogen (E2) in Hah et. al. (2011). Spearman’s Rho= 0.185, p < 2.2e-16. (**f**) Motifs enriched in TREs that have different amounts of RNA polymerase between TamS and TamR cells compared with TREs that have consistent levels.

### Distal enhancer activities correlate with tamoxifen resistance

To elucidate the mechanisms responsible for changes in gene transcription during the development of tamoxifen resistance, we sought to discover the location of promoters and active distal enhancers, collectively called transcriptional regulatory elements (TREs). Nascent transcription is a sensitive way to identify groups of active enhancers^11–14^, and results in enhancer predictions that are highly similar to the canonical active enhancer mark, acetylation of histone 3 at lysine 27 (H3K27ac)^12,13,25^. We used our dREG software package^13^ followed by a novel peak refinement step that identifies the regions between divergent paused RNA polymerase (see Methods; manuscript in preparation) to identify 39,753 TREs that were active in either the TamS or TamR MCF-7 lines. TREs discovered using dREG were highly enriched for other active enhancer and promoter marks in MCF-7 cells, especially H3K27ac (Supplementary Fig. 1a), as expected based on prior studies^11–13,25^. As an example, we selected a transcribed enhancer downstream of the *CCND1* gene for experimental validation using luciferase reporter gene assays, and confirmed luciferase activity in both B7TamS and G11TamR MCF-7 cells (Supplementary Fig. 1b and 1c).

We used the abundance of RNA polymerase recruited to each TRE as a proxy for its transcriptional activity in each MCF-7 subclone to identify differences in 1,452 TREs (812 increased and 640 decreased) (1% FDR, DESeq2) between TamS and TamR MCF-7 cells. Differentially transcribed TREs were frequently located near differentially expressed genes and undergo correlated transcriptional changes between the four MCF-7 subclones. *GREB1* and *PGR*, for example, are each located near several TREs, including both promoters (green) and enhancers (gray), which undergo changes between TamR and TamS MCF-7 cells that are similar in direction and magnitude to those of the primary transcription unit which encodes the mRNA (Fig. 2a and 2b). These results are consistent with a broad correlation between changes at distal TREs and protein coding promoters^11,24^.

We hypothesized that differential transcription at TREs reflects differences in the binding of specific transcription factors that coordinate changes between TamS and TamR lines. We identified 12 clusters of motifs enriched in TREs that are differentially active in the TamS and TamR lines (Bonferroni corrected p< 0.001; RTFBSDB^26^). The top scoring motif in this analysis corresponds to an estrogen response element (ERE), the canonical DNA binding sequence that recruits ERα to estrogen responsive enhancers (Fig. 2f). At least two of the top scoring motifs, those that were putatively bound by NFIA and HOX-family transcription factors (HOXC13 shown), bind a transcription factor that was itself differentially expressed in TamS and TamR MCF-7 cells (Fig. 2f), consistent with our expectation that transcriptional changes of a transcription factor elicit secondary effects on the activity of TREs, and downstream effects on gene transcription.

### ERα signaling remains functional in endocrine-resistant lines

*GREB1* and *PGR* play a critical role in ERα genomic activity in breast cancer cells^22,27^. Our observation that transcription of these ERα co-factors was lost in the resistant lines (Fig. 2a, 2b, 2c, and 2d) suggests that ERα signaling may be defective in the TamR cell lines. Consistent with this expectation, several analyses (i.e., the enrichment of ERα target genes and EREs, Fig. 1g and 1h) implicate global changes in the genomic actions of ERα during the development of tamoxifen resistance. However, these analyses are correlative and do not directly test the immediate responses to E2 in TamR and TamS lines.

To directly test the hypothesis that the genomic actions of ERα are substantially altered in the TamR lines, we treated B7^TamS^ and G11^TamR^ MCF-7 cells for 40 minutes with either E2 or tamoxifen, and monitored transcriptional changes using PRO-seq. RNA polymerase abundance increased sharply at ERα ChIP-seq peaks^28^ in B7^TamS^ MCF-7 cells (Fig. 3a top) in response to E2, but not in response to tamoxifen, in agreement with our prior work^11,29^. Although we observed a muted effect of E2 on enhancers in G11^TamR^ compared with B7^TamS^, increases in Pol II loading were observed in response to E2, but not tamoxifen (Fig. 3a bottom). These results demonstrate that E2 signaling pathway remains functional and able to affect gene transcription in a stimulus-dependent manner in TamR cells. We attribute the muted response in G11^TamR^ to a 2.44-fold reduction in the abundance of ERα mRNA in G11^TamR^ MCF-7 cells compared to the B7^TamS^ MCF-7 cells (Fig. 3b). This muted effect explains the enrichment in E2 target genes, as well as the ERE motif enrichment, between TamS and TamR lines shown in Fig. 1 and 2. Nevertheless, the genomic actions of E2-liganded ERα remain functional in TamR MCF-7 cells.

**Figure 3.**
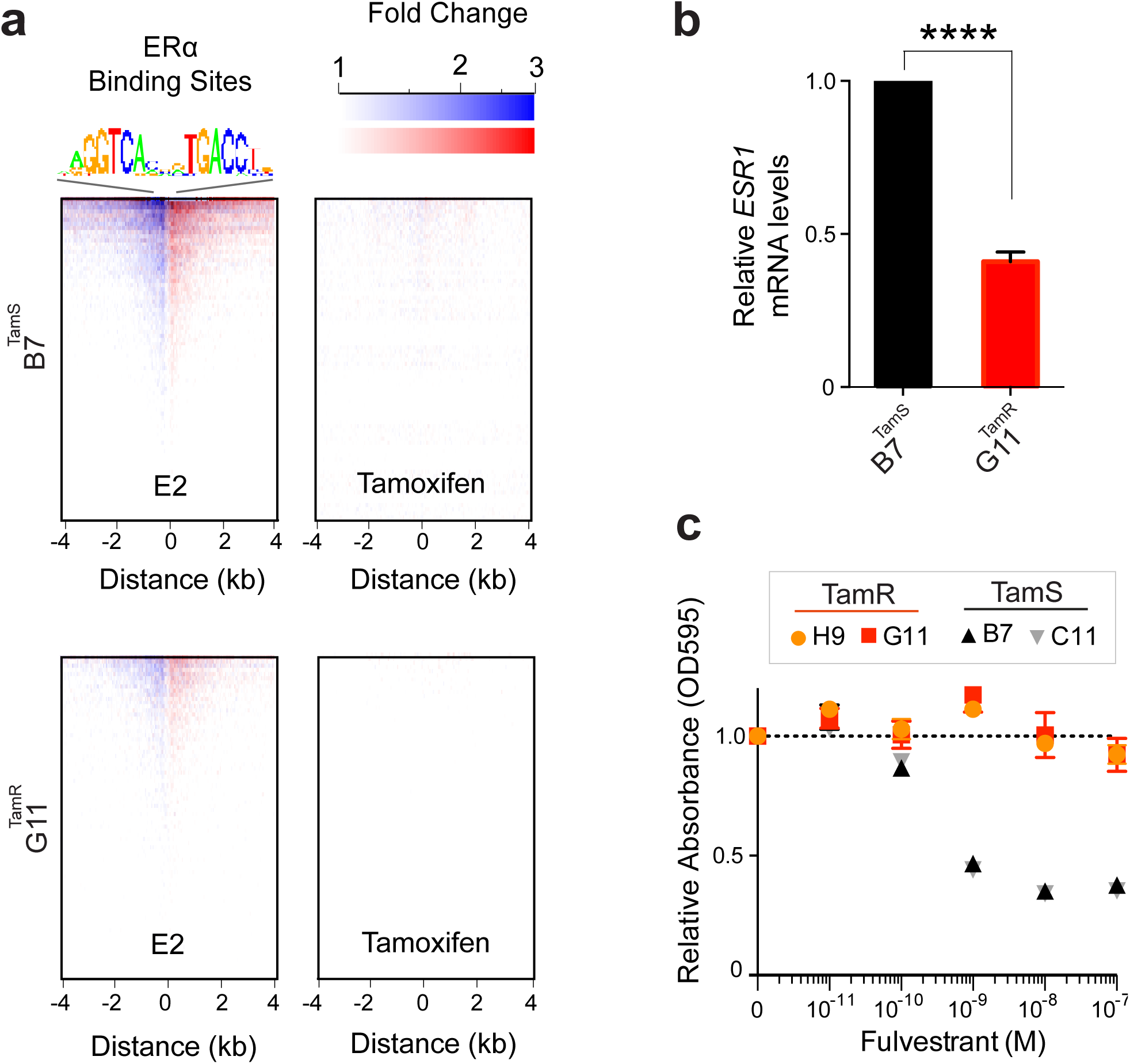
Tamoxifen resistant lines have functional ERα signaling. (**a**) Heatmap of changes in RNA polymerase abundance following 40 minutes of E2 or tamoxifen treatment near ERα bindings sites in B7^TamS^ and G11^TamR^ cells. (**b**) *ESR1* mRNA expression levels in B7^TamS^ and G11^TamR^ cells. Data are represented as mean ± SEM (n=3). **** p < 0.0001. (**c**) Cell viability of TamS and TamR cells upon treatment with 0 (vehicle; DMSO), 10^-11^, 10^-10^, 10^-9^, 10^-8^, or 10^-7^ M fulvestrant (ER degrader) for 4 days. Data are represented as mean ± SEM (n=3).

Given that E2 signaling remains functional, but muted in the TamR line, we next tested whether ERα was required for the growth of our tamoxifen-resistant cells. We found that the viability of both G11^TamR^ and H9^TamR^ MCF-7 cells was largely unaffected by treatment with the ER degrader, fulvesterant (Fig. 3c). Therefore, endocrine resistance in G11^TamR^ and H9^TamR^ MCF-7 cells appears to occur independently of ERα signaling, suggesting that these TamR lines are likely using an alternative pathway for cell survival and proliferation when grown in the presence of tamoxifen.

### GDNF is necessary and sufficient to confer endocrine resistance in MCF-7 cells

We next investigated pathways by which TamR lines may promote cell survival in the presence of endocrine therapies. Tyrosine kinase growth factor signaling pathways have been implicated in preclinical models of endocrine resistance^5,7,30^. RET is a cell surface receptor that elicits cell survival signals when bound by one of four RET ligands, GDNF, NRTN, ARTN, and PSPN^31^.One of these ligands, glial cell line-derived neurotrophic factor (GDNF), was among the most highly up-regulated genes in both G11^TamR^ and H9^TamR^ MCF-7 lines (Fig. 4a). We confirmed the transcriptional differences in *GDNF* between B7^TamS^ and G11^TamR^ MCF-7 cells using qPCR and found that GDNF mRNA levels were increased by ∼25 fold in the resistant line (Fig. 4b). Thus both *GDNF* transcription and mRNA abundance correlate with endocrine resistance in MCF-7 cells, suggesting that GDNF may contribute to the endocrine resistance phenotype.

**Figure 4.**
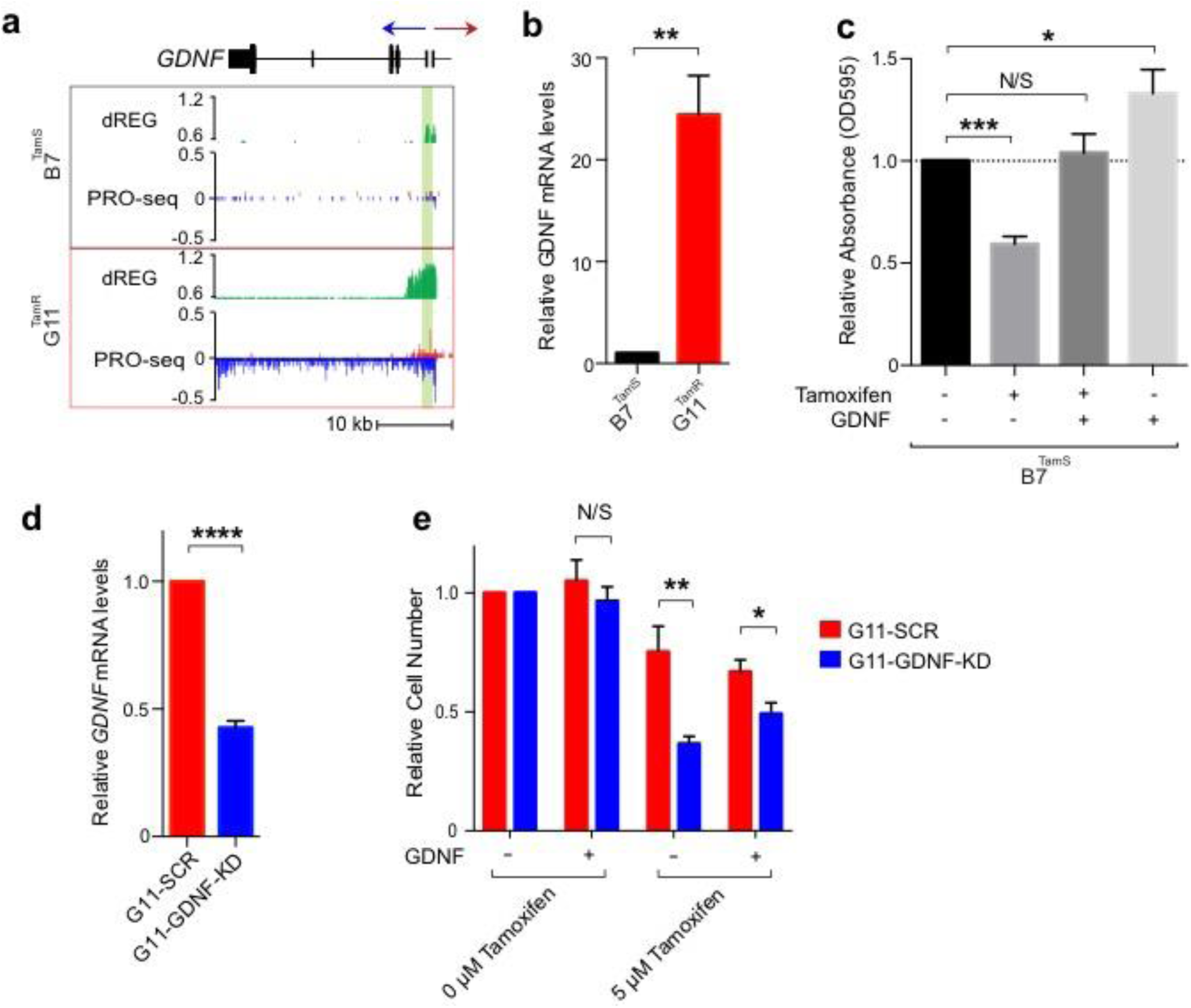
*GDNF* is responsible for tamoxifen resistance in MCF-7 cells. (**a**) Transcription near the *GDNF* locus in B7^TamS^ and G11^TamR^ cells. PRO-seq densities on sense strand and anti-sense strand are shown in red and blue, respectively. dREG scores are shown in green. The region near the GDNF promoter is shown in light green shading. Arrow indicates the direction of gene annotations. (**b**) *GDNF* mRNA expression levels in B7^TamS^ and G11^TamR^ cells. Data are represented as mean ± SEM (n=3). ** p < 0.005. (**c**) Cell viability of B7^TamS^ cells in the presence or absence of 10 ng/ml GDNF and/or 100 mM tamoxifen for 4 days. Data are represented as mean ± SEM (n=3). * p < 0.05, *** p < 0.0005. (**d**) *GDNF* mRNA expression levels in G11^TamR^ scrambled (SCR) and G11^TamR^ GDNF knockdown (GDNF-KD) cells. Data are represented as mean ± SEM (n=3). **** p < 0.0001. (**e**) Relative cell number of G11^TamR^ scrambled (SCR) and G11^TamR^ GDNF knockdown (GDNF-KD) cells after 4 days without or with 1 μM tamoxifen and/or 5 ng/ml GDNF treatment. Data are represented as mean ± SEM (n=9). * p < 0.05, ** p < 0.005.

We tested whether GDNF is casually involved in endocrine resistance by manipulating GNDF levels in our MCF-7 model. We first examined the effects of 10 ng/mL of recombinant GDNF protein on the growth of B7^TamS^ cells in the presence of antiestrogens. Remarkably, GDNF completely rescued B7^TamS^ MCF-7 cells when challenged with both tamoxifen (Fig. 4c) and fulvestrant (Supplementary Fig. 2a). Moreover, GDNF treatment without tamoxifen increased the proliferation rate of B7^TamS^ MCF-7 cells by ∼20% (Fig. 4c), suggesting that the growth pathways activated by GDNF can work independently of ERα. Next we tested whether GDNF was necessary to confer endocrine resistance in our model system by using short hairpin RNAs (shRNA) to knockdown GDNF in G11^TamR^ MCF-7 cells. Results show that GDNF depletion (GDNF-KD) reduced *GDNF* mRNA levels by 57.38% (Fig. 4d) and that these cells were significantly more sensitive to tamoxifen treatment than G11 cells transfected with a scrambled control (Fig. 4e). Moreover, endocrine resistance could be restored to GDNF-KD G11 cells by the addition of 5 ng/ mL recombinant GDNF protein (Fig. 4e), demonstrating that growth inhibition does not reflect an off-target effect of the *GDNF* shRNA. Taken together, these data demonstrate that *GDNF* plays a central and causal role in establishing endocrine resistance in G11^TamR^ MCF-7 cells.

### Endocrine-sensitive ER+ breast cancer cells express RET transmembrane receptors

Having shown that *GDNF* expression promotes endocrine resistance in our MCF-7 cell model, we asked whether *GDNF* promotes resistance in patients as well. Increases in the expression of RET tyrosine kinase or its co-receptor GFRα1 are thought to be involved in endocrine resistance^5–7^. However, RET is itself transcriptionally activated by ERα and is highly abundant in endocrine sensitive ER+ breast cancer cell models^24^. Analysis of mRNA-seq data from 1,177 primary breast cancers in the cancer genome atlas (TCGA) revealed that the RET mRNA expression level was highest in ER+ breast cancer and correlates positively with the expression level of *ESR1* (ERα) (Spearman’s *ρ* = 0.51, *p* < 2.2e-16; Fig. 5a), suggesting that it is a direct transcriptional target of ERα *in vivo* as well. *GFRA1* mRNA encodes the GDNF co-receptor, GFRα1, and, together with RET, activates RET-ligand signaling. Further analysis of the mRNAseq data set found that *GFRA1* is also strongly correlated with *ESR1* mRNA in breast cancers (Spearman’s *ρ* = 0.67, *p* < 2.2e-16; Supplementary Fig. 3a), suggesting that it is also a direct target of E2 signaling. In our MCF-7 endocrine resistance model, *GFRA1* transcription is 5-fold higher in TamS MCF-7 cells compared to TamR lines and *RET* transcription is not significantly different (Fig. 5b and 5c), demonstrating that neither factor is overexpressed in TamR MCF-7 cells. Since both RET and GFRA1 are naturally high in ER+ breast cancer cells, and since high expression of these factors appears to be established in part by ERα, there must be other causes of endocrine resistance, both in cell models and *in vivo*.

**Figure 5.**
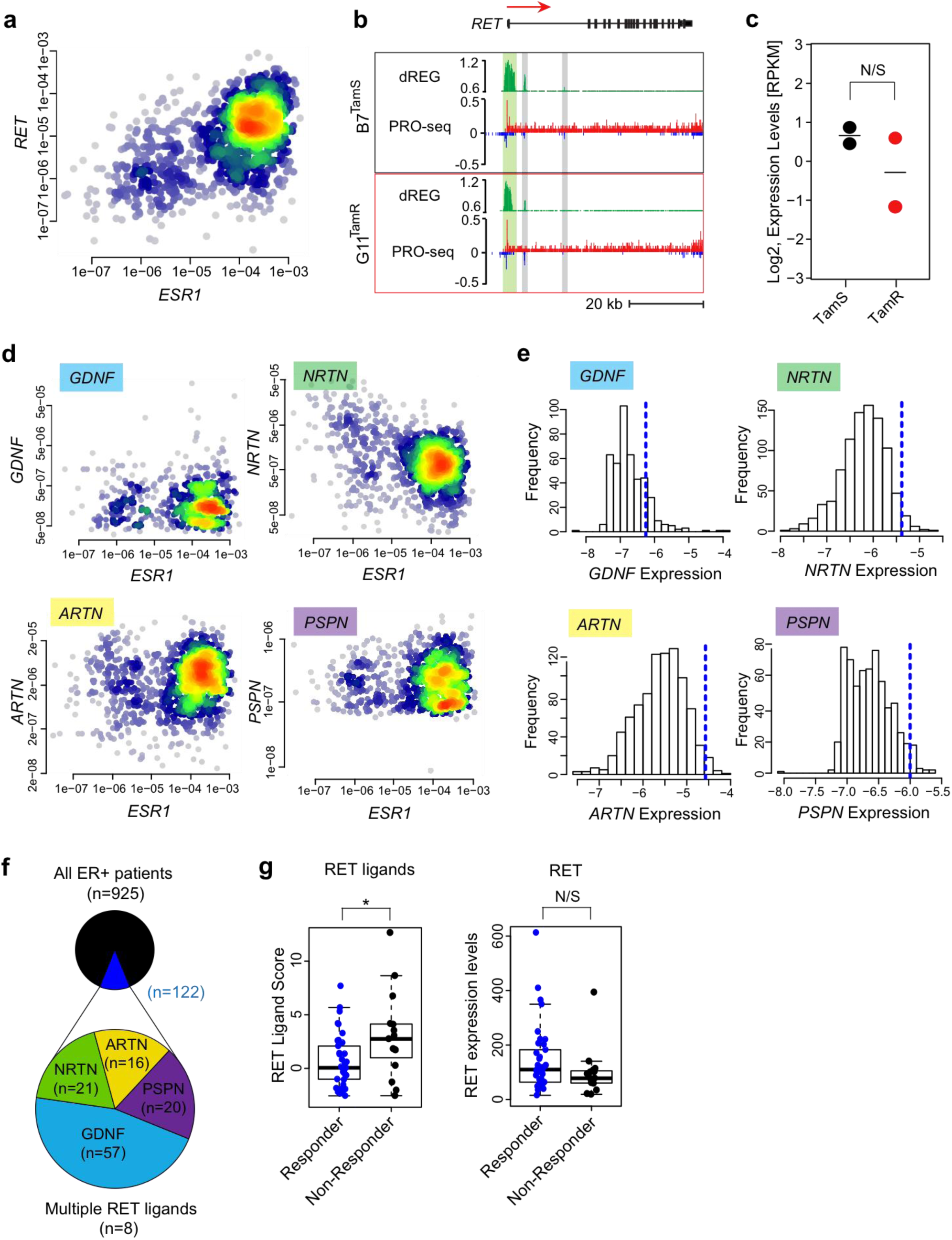
Expression of RET ligands contributes to endocrine resistance. (**a**) Density scatterplot showing *RET* and *ESR1* expression in mRNA-seq data from 1,177 primary breast cancer models in the cancer genome atlas (TCGA). Spearman’s ρ = 0.51, *p* = 1.2e-60. (**b**) Transcription near the *RET* locus in B7^TamS^ and G11^TamR^ cells. PRO-seq densities on sense strand and anti-sense strand are shown in red and blue, respectively. dREG scores are shown in green. Enhancers and promoters are shown in grey and light green shading, respectively. Arrow indicates the directional movement of transcribed genes. (**c**) Dot plot shows *RET* transcription levels in TamS and TamR MCF-7 cells. (**d**) Density scatterplots show the expression of RET ligands (*GDNF*, *NRTN*, *ARTN*, and *PSPN*) versus *ESR1* based on mRNAseq data from 1,177 primary breast cancers. (**e**) RET ligand expression distribution in ER+ breast cancers. The dotted blue line represents 2.5 times the range between the 25^th^ and 50^th^ percentile. (**f**) Fraction of ER+ breast cancers (n = 925) with at least one RET ligand exceeding the threshold shown in panel E (shown in dark blue, n = 122). Among the 4 RET ligands, GDNF was the most highly expressed (n = 60). (**g**) Boxplots show RET ligands score and RET expression levels in patients that respond or do not respond to aromatase inhibitor letrozole. * *p* = 0.016.

### ER+ breast cancer cells and primary breast cancers that are sensitive to endocrine therapy lack GDNF to initiate resistance

Our finding that recombinant GDNF was sufficient for endocrine resistance in B7^TamS^ MCF-7 cells demonstrates that GDNF is a key limiting factor, the absence of which prevents TamS cells from developing a resistant phenotype. To extend this hypothesis to primary breast cancers, we sought to determine whether GDNF expression is normally low, such that it might limit RET pathway activation in most ER+ breast cancers. Indeed, GDNF expression was detectible in only 565 of 1,177 primary breast cancers (48%) analyzed by TCGA (Supplementary Fig. 3b). In principal, RET signaling may be activated by any of the four RET ligands (GDNF, NRTN, ARTN, and PSPN). However, only low levels of NRTN, ARTN, or their co-receptors were detected in primary breast tumors (Fig. 5d, 5e, and Supplementary Fig. 3b). Thus, we conclude that RET ligand expression is low compared with that of cell surface receptors, especially RET and GFRα1, which are activated in part by ERα. This contrast between RET receptors and ligands supports a model in which the RET signaling pathway is ‘poised’ for endocrine resistance by expression of the receptors and that limiting levels of *GDNF* expression, or possibly other RET ligands, would ensure endocrine sensitivity in most tumors.

Next, we investigated whether high RET ligand expression in a subset of ER+ tumors may explain some cases of endocrine resistance. A careful examination of the GDNF expression distribution in TCGA breast cancers revealed a long tail, indicating high GDNF expression in a subset of cases in the TCGA dataset (Fig. 5e). Our hypothesis that GDNF expression limits RET-dependent endocrine resistance implies that these GDNF-high samples should be prone to endocrine resistance. We devised a simple non-parametric computational approach, which we call the ‘outlier score’, to quantify the degree to which GDNF is highly expressed based on the symmetry of the empirical probability density function (see Methods; Fig. 5e, blue line). Based on this score, we conservatively estimate that, of 925 ER+ breast cancer patients in the TCGA dataset, 122 have high expression of at least one of the RET ligands (13%), 57 of which had high levels of GDNF (Fig. 5f).

If RET ligands are the limiting factor for endocrine resistance, as we propose here, cases included in this long distribution tail are those that are more likely to be resistant to endocrine therapies. To test this hypothesis, we analyzed expression microarray data collected prospectively by biopsies of patients that either respond, or do not respond, to the aromatase inhibitor letrozole^32^. A score comprised of the sum of the outlier scores from all four RET ligands is significantly higher in patients that do not respond to letrozole treatment (p= 0.016, one-sided Wilcoxon rank sum test; Fig. 5g). By contrast, there is no significant difference in RET expression in patients who respond or who do not respond to letrozole. These results suggest that RET ligand expression, but not RET itself, explain the differences in response to letrozole in this cohort of patients.

## Discussion

In this study, we have used genomic tools to dissect how the GDNF-RET signaling pathway becomes activated in breast cancer cells to promote resistance to endocrine therapies. Systematic experimental manipulation of GDNF expression in TamS and TamR cell lines build on work described in previous studies^5–8^ by providing the strongest support yet for this pathway playing a causal role in endocrine resistance in MCF-7 cells. Furthermore, analysis of clinical data points toward a model in which *RET* and *GFRA1* are actively transcribed in both endocrine sensitive MCF-7 cells and primary tumors, awaiting RET ligands to initiate resistance to endocrine therapies. This is, to our knowledge, the first study to suggest that expression of RET ligands themselves (including GDNF, ARTN, NRTN, and PSPN) are responsible for RET-mediated endocrine resistance. Overall, our study provides insights into how the RET signaling pathway become activated in ER+ breast cancers.

We are the first to propose that RET-mediated endocrine resistance occurs when ER+ breast cancer cells express the RET ligand GDNF. Work on the RET signaling pathway in endocrine resistance has largely focused on amplifications or increases in the expression of RET or its co-receptor GFRα1 in resistance to aromatase inhibitors^6,7^. However, RET expression is not significantly different in a cohort of patients resistant to the aromatase inhibitor letrozole (Fig. 5g), suggesting that other mechanisms may occur more commonly in patients than differences in the expression of RET itself. Indeed, we find that expression of RET and GFRα1 are both highest in ER+ breast cancers, likely because of direct transcriptional activation of both genes by E2/ ERα (Fig. 5a and Supplementary Fig. 3a). Thus, we propose that ER+ breast cancer cells are intrinsically ‘poised’ for RET-mediated endocrine resistance by the activation of RET cell-surface receptors, but lack expression of the ligand GDNF.

Based on these findings, we hypothesize that increased expression of any one of the four RET ligands, GDNF, ARTN, NRTN, or PSPN confers endocrine resistance on cells expressing the RET receptor. In support of this model, we demonstrate that the scoring system we used, based on RET ligand overexpression in tumors, clearly separates breast cancer patients that respond to letrozole from those who do not (Fig. 5g). Several findings also strongly support the involvement of GDNF in endocrine resistance in our MCF-7 model, most notably the observations that GDNF rescues B7^TamS^ lines and that GDNF knockdown in G11 cells restores sensitivity to tamoxifen (Fig. 4e). These observations are also supported by existing studies showing that another RET ligand, ARTN, contributes to tamoxifen resistance in MCF-7 cells^33^, extending and supporting the findings reported here. However, there is one RET ligand that is notably an outlier. PSPN does not appear to have any predictive value in patients, and thus may not play the same role in resistance as the other three RET ligands. This may reflect the extremely low expression of its co-receptor, *GFRA4*, in primary breast cancers (Supplementary Fig. 3b), preventing PSPN from having much effect on breast cancer cells. Taken together, these findings suggest that RET ligand expression, especially GDNF, ARTN, and NRTN, explain endocrine resistance in many cases.

A major question that remains unclear and of primary importance following our study is how RET ligand expression becomes activated in primary tumors. The abundance of GDNF mRNA appears to be extremely low in primary breast tumors analyzed by TCGA (Fig. 5d, 5e, and Supplementary Fig. 3b), which were in most cases collected before therapeutic intervention^34,35^. Notably, GDNF is not natively expressed in ER+ TamS MCF-7 cells but rather becomes activated following extended GDNF treatments. This may suggest that GDNF expression is initiated in tumors by another stimulus-dependent pathway or introduced by another cell type in the tumor microenvironment. Consistent with this, GDNF expression in tumors may require pro-inflammatory cytokines, such as tumor necrosis factor alpha (TNFα), to be transcribed in breast cancer cells^23^. This finding may link poor survival outcomes in pro-inflammatory tumors^36,37^ with GDNF-RET-mediated resistance to endocrine therapy.

Taken together, results reported in this study implicate RET ligands, including GDNF, as the primary determinant of endocrine resistance in both MCF-7 cells and patient samples (Fig. 6). Clinical studies targeting larger cohorts of patients beginning endocrine therapies will be required to fully validate our proposed mechanism of endocrine resistance.

**Figure 6.**
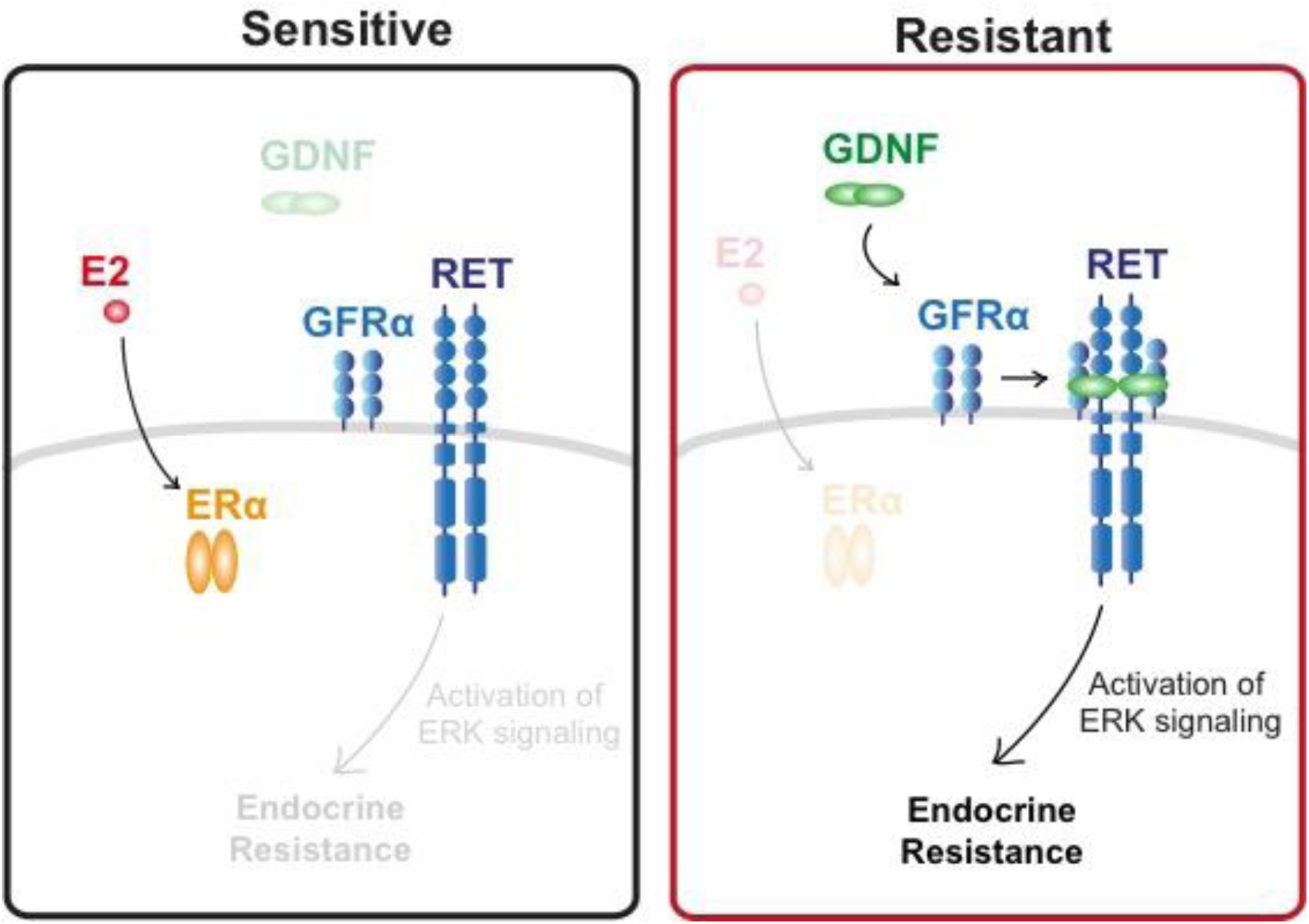
Schematic diagram of RET activation in endocrine sensitive and resistant tumors. Both endocrine sensitive and resistant breast cancer cells express all components of the RET signaling pathway, but endocrine sensitive breast cancer cells lack GDNF to initiate the resistance pathway.

## Methods

### Cell lines and cell culture

Tamoxifen-sensitive (TamS; B7^TamS^ and C11^TamS^) and resistant (TamR; G11^TamR^ and H9^TamR^) MCF-7 cells^9^ were a gift from Dr. Joshua LaBaer. TamS cells were grown in Dulbecco’s Modified Eagle Medium supplemented with 5% fetal bovine serum and 1% Penicillin Streptomycin, and TamR cells were grown in the same media supplemented with 1 μM tamoxifen. Tamoxifen used throughout in this paper is (Z)-4-Hydroxytamoxifen (4-OHT; Sigma-Aldrich; Cat# H7904).

### Cell viability assay

Briefly, 5 x 10^3^ TamS and TamR cells were grown in 24-well TC-treated plates in their specific culture media. After allowing the cells to adhere to the plate for 24 hours, they were rinsed with PBS three times to remove any residual tamoxifen. The cells were treated with either increasing doses of tamoxifen (0 (vehicle control; EtOH), 10^-11^, 10^-10^, 10^-9^, 10^-8^, or 10-^7^ M).

For setting up the rescue experiment with GDNF (PeproTech; Cat# 450-10), 5 x 10^3^ B7^TamS^ cells were grown in 24-well TC-treated plates in their specific culture media. After allowing the cells to adhere to the plate for 24 hours, they were treated with either EtOH (vehicle), 10^-7^ M tamoxifen, 10^-7^ M tamoxifen and 10 ng/mL GDNF, or 10 ng/mL GDNF treatment. The same set up was performed for 10^-7^ M treatment of fulvestrant and using DMSO (vehicle) as a control.

After four days of endocrine treatment, cells were fixed with 4% araformaldehyde and stained with 0.5% crystal violet solution made in 25% methanol. After washing away non-specific crystal violet stain with PBS, we took pictures of each plate and the crystal violet stain from the fixed cells was removed using 10% acetic acid. The absorbance was measured using the Tecan plate reader at OD595nm. Samples were normalized to the untreated control. Three biological replicates were performed and data are represented as mean ± SEM.

### Cell culture set up and nuclei isolation

TamS and TamR lines were grown in 150mm TC-treated culture dishes in their respective normal culture media. Cells were rinsed with PBS at least three times 24 hours after plating. Both the TamS and TamR cells were grown in Dulbecco’s Modified Eagle Medium supplemented with 5% fetal bovine serum and 1% Penicillin Streptomycin for an additional three days until ∼80% confluency in the absence of tamoxifen, in order to measure the difference between TamS and TamR cells pre-treatment.

Nuclei were isolated as described previously ^38^. Briefly, cells were rinsed three times with ice-cold PBS and lysed using lysis buffer (10 mM Tris-HCl pH 7.4, 2 mM MgCl_2_, 3 mM CaCl_2_, 0.5% NP-40, 10% Glycerol, 1 mM DTT, 1X PIC (Roche; Cat# 11836153001), and 1 μl/10 mL SUPERase-In (ThermoFisher; Cat# AM2694) dissolved in DEPC water). Cells were homogenized by gently pipetting at least 30 times and the nuclei were harvested by centrifugation at 1000 *g* for five minutes at 4°C. The isolated nuclei were washed twice with lysis buffer and were resuspended in 100 μL freezing buffer (50 mM Tris HCl pH 8.3, 5 mM MgCl_2_, 40% Glycerol, 0.1 mM EDTA pH 8.0, and 4 U/mL SUPERase-In). The isolated nuclei were used for nuclear run-on and precision nuclear run-on sequencing (PRO-seq) library preparation.

### Nuclear run-on and PRO-seq library preparation

Nuclear run-on experiments were performed according to the methods described previously ^17,18^. 1x10^7^ nuclei in 100 μL freezing buffer were mixed with 100 μL of 2x nuclear run-on buffer (10 mM Tris-HCl pH 8.0, 5 mM MgCl2, 1 mM DTT, 300 mM KCl, 50 μM biotin-11-ATP (Perkin Elmer; Cat# NEL544001EA), 50 μM biotin-11-GTP (Perkin Elmer; Cat# NEL545001EA), 50 μM biotin-11-CTP (Perkin Elmer Cat# NEL542001EA), 50 μM biotin-11-UTP (Perkin Elmer; Cat# NEL543001EA), 0.4 units/μL SUPERase In RNase Inhibitor (Life Technologies; Cat# AM2694), 1% Sarkosyl (Fisher Scientific; Cat# AC612075000). The mixture was incubated at 37 °C for five minutes. The biotin run-on reaction was stopped using Trizol (Life Technolgies; Cat# 15596-026), Trizol LS (Life Technologies; Cat# 10296-010) and pelleted. The use of GlycoBlue (Ambion; Cat# AM9515) is recommended for higher pellet yield. RNA pellets were re-dissolved in DEPC water and denatured in 65 °C for 40 seconds and hydrolyzed in 0.2 N NaOH on ice for 10 minutes to have a hydrolyzed RNA length with that range ideally of 40 to 100 nts. Bead binding (NEB; Cat# S1421S) was performed to pull down nascent RNAs followed by 3’ RNA adaptor ligation (NEB; Cat# M0204L). Another bead binding was performed followed by 5’ de-capping using RppH (NEB; Cat# M0356S). 5’ phosphorylation was performed followed by 5’ adaptor ligation. The last bead binding was performed before generation of cDNA by reverse transcription. PRO-seq libraries were prepared according to manufacturers’ protocol (Illumina) and were sequenced using the Illumina NextSeq500 sequencing.

### Mapping of PRO-seq sequencing reads

PRO-seq reads failing Illumina quality filters were removed. Adapters were trimmed from the 3’ end of remaining reads using cutadapt with a 10% error rate ^39^. Reads were mapped with BWA^40^ to the human reference genome (hg19) and a single copy of the Pol I ribosomal RNA transcription unit (GenBank ID# U13369.1). The location of the RNA polymerase active site was represented by a single base that denotes the 3’ end of the nascent RNA, which corresponded to the position on the 5’ end of each sequenced read. Mapped reads were normalized to reads per kilobase per million mapped (RPKM) and converted to bigWig format using BedTools^41^ and the bedGraphToBigWig program in the Kent Source software package^42^. Downstream data analysis was preformed using the bigWig software package, available from: https://github.com/andrelmartins/bigWig. All data processing and visualization was done in the R statistical environment^43^.

### Identification of active enhancers and promoters using dREG-HD

We identified TREs using dREG ^13^. Data collected from all four cell lines (TamR and TamS MCF-7 cells) was combined to increase statistical power for the discovery of a superset of TREs active during any of the conditions examined.

The precise coordinates of TREs were refined using a strategy that we termed dREGHD (available at https://github.com/Danko-Lab/dREG.HD; manuscript in preparation). Briefly, dREG-HD uses an epsilon-support vector regression (SVR) with a Gaussian kernel to map the distribution of PRO-seq reads to DNase-I signal intensities. Training was conducted on randomly chosen positions within dREG peaks in K562 cells (GEO ID# GSM1480327) extended by 200bp on either side. We selected the optimal set of features based on maximizing the Pearson correlation coefficient between the imputed and experimental DNase-I signal intensity over an independent validation set. Before DNase-I imputation, PRO-seq data was preprocessed by normalizing read counts to the sequencing depth and scaled such that the maximum value was within the 90^th^ percentile of the training examples. To identify peaks, we smoothed the imputed DNase-I signal using a cubic spline and identified local maxima. We tuned the performance of the peak by empirically optimizing two free parameters that control the (1) smoothness of spline curve fitting, and (2) a threshold level on the intensity of the imputed DNase-I signal. Parameters were optimized to achieve <10% false discovery rates on a K562 training dataset by a grid optimization over free parameters. We tested the optimized dREG-HD model (including both DNase-I imputation and peak calling) a GRO-seq dataset completely held out from model training and parameter optimization in on GM12878 lymphoblastoid cell lines (GSM1480326). Testing verified that dREG-HD identified transcribed DNase-I hypersensitive sites with 82% sensitivity at a 10% false discovery rate.

Additional genomic data in MCF-7 cells generated by the ENCODE project was downloaded from Gene Expression Omnibus. TREs discovered using dREG-HD were compared with ChIP-seq for H3K27ac and H3K4me3 (accession numbers: GSM945854 and GSM945269) and DNase-1 hypersensitivity (GSM945854).

### Differential expression analysis (DESeq2)

We compared treatment conditions or cell lines using gene annotations (GENCODE v19). We counted reads in the interval between 1,000 bp downstream of the annotated transcription start site to the end of the gene for comparisons between TamS and TamR cell clones. To quantify transcription at enhancers, we counted reads on both strands in the window covered by each dREG-HD site. Differential expression analysis was conducted using deSeq2 ^19^ and differentially expressed genes were defined as those with a false discovery rate (FDR) less than 0.01.

### Motif enrichment analysis

Motif enrichment analyses were completed using the default set of 1,964 human motifs in RTFBSDB^26^ clustered into 622 maximally distinct DNA binding specificities (see ref Wang et. al. (2016)). We selected the motif to represent each cluster with canonical transcription factors that were most highly transcribed in MCF-7 cells. We fixed the motif cutoff log odds ratio of 7.5 (log *e*) in a sequence compared with a third-order Markov model as background. We identified motifs enriched in dREG-HD TREs that change transcription abundance between two conditions using Fisher’s exact test with a Bonferroni correction for multiple hypothesis testing. TREs were compared to a background set of >1,500 GC-content matched TREs that do not change transcription levels (<0.25 absolute difference in magnitude (log-2 scale) and *p* > 0.2) using the enrichmentTest function in RTFBSDB^26^.

### TCGA data analysis

Processed and normalized breast cancer RNA-seq data was downloaded from the International Cancer Genome Consortium (ICGC) data portal website (https://dcc.icgc.org). Data profiling each gene was extracted using shell scripts. Processing and visualization was done in R.

### Letrozole microarray reanalysis

We reanalyzed Affymetrix U133A microarray data profiling mammary tumor biopsies before and after treatment with letrozole^32^. Miller et. al. (2012) collected data from mammary tumor biopsies prior to letorozle treatment, 10-14 days following the start of treatment, and 90 days following the start of treatment. Samples were annotated as a “responder” (i.e., responds to letrozole treatment), a “non-responder” (i.e., no benefit from letrozole treatment), or “not assessable” (i.e., unknown). The Series Matrix Files were downloaded from Gene Expression Omnibus (GSE20181) and each gene of interest was extracted and processed into a text file. We used the following Affymetrix ID numbers 221359_at, 210683_at, 210237_at, 221373_x_at, 211421_s_at, and 201694_s_at to represent *GDNF*, *NRTN*, *ARTN*, *PSPN*, *RET*, and *EGR1*, respectively. We found no evidence of differences in RET or RET ligand expression across the three time points, and we therefore used the average expression of each RET ligand in each sample when comparing between responsive and non-responsive patients in order to decrease assay noise.

Outlier scores were designed to score the degree to which each sample fell within the tail of the distribution representing high expression levels of each RET ligand (as shown in Fig. 4E). Because endocrine resistance could, in principal, be caused either by high expression of any individual RET ligand on its own, or by moderately high expression of multiple RET ligands in combination, we devised a data transformation and sum approach to score the degree to which all four of the RET ligands were highly expressed in each sample. In our data transformation, expression levels were centered by the median value and scaled based on the lower tail of the expression distribution (between quartile 0 and 50). This approach is similar in concept to a Z-score transform, but uses the lower tail to estimate the variance in order to avoid having high expression levels, which we hypothesize here may contribute to endocrine resistance, from contributing to the denominator used to standardize the distribution of each RET ligand. After transforming scores from all four RET ligands separately, we took the sum of the scores to represent our final ‘outlier score’. Because our hypothesis specifically predicted an increase in the RET ligand score to correlate with letrozole resistance, and because the number of patients was small, we designed the analysis to use a one-tailed Wilcoxon rank sum test. However, in practice, using a two-tailed Wilcoxon rank sum test did not change the results of our analysis. Data was processed and visualization was completed using R.

### RNA isolation and quantitative real-time PCR

RNA was purified using an RNeasy Kit (Qiagen; Cat# 74104) and 1μg of purified RNA was reverse-transcribed using a High Capacity RNA-to-cDNA kit (Applied Biosystems; Cat# 4387406) according to the manufacturers’ protocols. Real-time quantitative PCR analysis was performed using the following primers: *ACTB* Forward (5’-CCAACCGCGAGAAGATGA-3’) and Reverse (5’-CCAGAGGCGTACAGGGATAG-3’); *PGR* Forward (5’-GTCAGGCTGGCATGGTCCTT-3’) and Reverse (5’-GCTGTGGGAGAGCAACAGCA-3’); *GREB1* Forward (5’-GTGGTAGCCGAGTGGACAAT-3’) and Reverse (5’-ATTTGTTTCCAGCCCTCCTT-3’) ^44^; *GDNF* Forward (5’-TCTGGGCTATGAAACCAAGGA-3’) and Reverse (5’-GTCTCAGCTGCATCGCAAGA-3’)^45^; and Power SYBR Green PCR Master Mix (Applied Bioystems; Cat#4367659). The samples were normalized to β-actin. At least three biological replicates were performed and data are presented as mean ± SEM. All statistical analyses for qPCR were performed using GraphPad Prism. Groups were compared using a two-tailed unpaired Student’s t-test.

### Generation of GDNF knockdown G11 cells

*GDNF* expression was stably knocked down in G11^TamR^ cells by transduction with lentivirus expressing either a shRNA scrambled control or *GDNF* shRNA. Mission shRNA lentivirus plasmids for control shRNA (Cat# SHC002) and *GDNF* shRNA (Cat# SHCLND-NM_000514) from Sigma-Aldrich were used. Specifically, 1.5 μg pLKO.1 shRNA plasmid (Addgene; Plasmid #1864), 0.5 μg psPAX2 packaging plasmid (Addgene; Plasmid #12260), and 0.25 μg pMD2.G envelope plasmid were used for packaging (Addgene; Plasmid #12259). The lentiviruses were generated and transduced according to the manufacturer’s instructions (Sigma-Aldrich). Clones were selected in 2 μg/ml of puromycin.

### Cell proliferation assay

Approximately 1x10^6^ G11-scrambled (G11-SCR) and G11-GDNF-knockdown (G11-GDNF-KD) cells were plated in T25 TC-flask. The cells were grown in either 0, 1 or 10 μM tamoxifen in the presence or absence of 5 ng/mL GDNF for 7 days. The cell number was counted for quantification and was normalized to the untreated group. Three biological replicates were performed.

### Statistical analysis

Statistical parameters include the exact number of biological replicates (n), standard error of the mean (mean ± SEM), and statistical significance are reported in the figure legends. Data are reported statistically significant when p < 0.05 by two-tailed Student’s t-test. In figures, asterisks and pound signs denote statistical significance as calculated by Student’s t-test. Specific p-values are indicated in the figure legends. Statistical analysis was performed using GraphPad PRISM 6.

## Acknowledgements

We thank X. Yao, L. Lan, as well as all members of the Danko and Coonrod labs for valuable insights and discussions. We also thank G. Leiman for edits and J. Lewis for input on the manuscript draft. Work in this publication was supported by NHGRI (National Human Genome Research Institute) grants from the US National Institutes of Health under award number R01 HG009309-01 to CGD. The content is solely the responsibility of the authors and does not necessarily represent the official views of the US National Institutes of Health.

## Author contributions

The project was conceived by CGD, SAC, and SH. All cell culture and molecular experiments were done by SH, HZ, and LJA. PRO-seq experiments were conducted by EJR and SH. Genomic data was analyzed by CGD and SH. Data collection, experiments, and analysis was supervised jointly by CGD and SAC. The paper was written by SH, CGD, and SAC with input from all authors.

## Data availability

Raw data files for the PRO-seq analysis have been deposited in Gene Expression Omnibus under Accession Number GSE93229. This study can be viewed before official release at: https://www.ncbi.nlm.nih.gov/geo/query/acc.cgi?token=cpmbkeuyxxmjdaj&acc=GSE93229. Software and scripts used in all analyses are publicly available without restriction on GitHub at https://github.com/Danko-Lab/mcf7tamres. At the time of submission, the most recent commit was version number: 855156ad07c042c88089cb4f31bf9d544487a1b2.

